# TimeCycle: Topology Inspired MEthod for the Detection of Cycling Transcripts in Circadian Time-Series Data

**DOI:** 10.1101/2020.11.19.389981

**Authors:** Elan Ness-Cohn, Rosemary Braun

## Abstract

**Motivation:** The circadian rhythm drives the oscillatory expression of thousands of genes across all tissues. The recent revolution in high-throughput transcriptomics, coupled with the significant implications of the circadian clock for human health, has sparked an interest in circadian profiling studies to discover genes under circadian control.

**Result:** We present TimeCycle: a topology-based rhythm detection method designed to identify cycling transcripts. For a given time-series, the method reconstructs the state space using time-delay embedding, a data transformation technique from dynamical systems theory. In the embedded space, Takens’ theorem proves that the dynamics of a rhythmic signal will exhibit circular patterns. The degree of circularity of the embedding is calculated as a persistence score using persistent homology, an algebraic method for discerning the topological features of data. By comparing the persistence scores to a bootstrapped null distribution, cycling genes are identified. Results in both synthetic and biological data highlight TimeCycle’s ability to identify cycling genes across a range of sampling schemes, number of replicates, and missing data. Comparison to competing methods highlights their relative strengths, providing guidance as to the optimal choice of cycling detection method.

**Availability and Implementation:** A fully documented open-source R package implementing TimeCycle is available at: https://nesscoder.github.io/TimeCycle/.

## Introduction

Circadian rhythms—physiological, behavioral, and metabolic oscillations with an approximate 24-h period—are controlled by an evolutionarily conserved set of core clock genes operating at the transcriptional and protein level. Entrained by Zeitgebers (external environmental stimuli such as light, temperature, and food) that modulate time–of–day specific functions, the circadian clock orchestrates a multitude of cellular processes, including nearly half of genes across all tissues [1]. Although various epidemiological studies have established significant links between circadian rhythms and human health [1–10], the underlying biological mechanisms coupling these phenomena remain poorly understood.

Facilitated by the development of high-throughput assays, researchers can now interrogate biological mechanisms at the molecular level by analyzing transcriptomic time-series data to identify genes under circadian control. This capability presents researchers with new analytical challenges of how best to reliably extract rhythmic signals from transcriptomic time-series data [11]. First, experimental costs constrain the frequency and length of sampling, requiring conclusions to be made from sparse or short time-series measurements. Second, circadian expression profiles often do not follow precise sinusoidal waveforms, but exhibit asymmetries, sharp peaks, additive trends, and noisy fluctuations [12–16]. To address these challenges, a variety of cycling detection methods have been proposed [12, 13, 15, 17–25]. However, bench-marking studies indicate that no method is universally optimal, and different algorithms may yield conflicting results on the same data [16, 26–29].

Early cycling detection used parametric approaches employing auto-correlation, curve fitting, and Fourier analysis to decompose expression patterns into harmonic components of varying amplitude and phase [17–19]. These methods could successfully identify genes with relatively symmetric waveform shapes, but had poor performance on asymmetric and sharply peaked waveforms [12, 13]. To address these shortcomings, a range of non-parametric methods were developed [13,15,22–25]. While non-parametric methods sacrifice statistical power relative to parametric methods [28], they are applicable to a broader range of cycling waveforms and generally outperform parametric methods [22, 23, 25, 30].

Nevertheless, limitations remain in current methods [11, 16, 28, 31]. A common approach is to compare the observed gene expression time-series to a user-defined set of reference waveforms (e.g. oscillations with different degrees of sawtooth asymmetry) to calculate a periodicity score (e.g. via the Kendall *τ* rank correlation). Such methods may fail to detect cyclic patterns that do not fall into the predetermined profiles of reference signals, effectively limiting the scope of discovery. Moreover, these methods typically assess the statistical significance relative to a null model of a randomized time-series. Since randomized time-series may jump from low to high gene expression faster than biological translation and degradation processes allow, these methods may produce unrealistic null models and misleading significance tests.

In addition to methodological hurdles, there are also considerations of various methods’ abilities to handle replicates, uneven sampling, missing data, and computational efficiency. In practice, the ability of methods to adequately handle these features directly affects flexibility in experimental design. For instance, a method that can accommodate uneven sampling can allow for dense sampling at times of interest, with sparser sampling at other times. Because missing data often occur as a result of sequencing errors with greater likelihood as sample size increases [32], researchers benefit from algorithms that can adequately handle missingness. Finally, computational efficiency allows for data set sizes to grow while still processing the data in a reasonable amount of time.

Results from dynamical systems theory and toplogogical data analysis provide alternative strategies to overcome the limitations of curve-fitting and template-based analysis methods. The recent “SW1PerS” method [24] uses time–delay embedding [33] to transform sliding windows of the time-series into a high-dimensional point cloud, and quantifies the circularity of the transformed signal as a measure of its periodicity. However, because the rhythmicity scores do not follow a well-defined distribution and the computational complexity of the algorithm precludes permutation tests, SW1PerS does not provide a *p*-value testing whether a gene is cycling.

To address these challenges, we introduce TimeCycle: a nonparametric, template–free algorithm based on topological data analysis with a bootstrapping procedure for statistical inference of cycling genes. Results demonstrate that TimeCycle reliably discriminates cycling and non-cycling profiles in synthetic data and reproducibly detects known circadian genes in experimental data. Below we describe the method, illustrate its application to multiple data sets, and provide a comparison to several competing methods.

## Results

TimeCycle is a method for classifying and quantifying cyclic patterns of gene expression in transciptomic time-series data. Application to both synthetic and experimental data demonstrate TimeCycle’s ability to efficiently discriminate cycling genes from non-cycling genes across a range of sampling schemes, number of replicates, and missing data.

### The TimeCycle Method

TimeCycle’s basic framework consists of a rescaling/normalization step, reconstruction of the state space via time-delay embedding, isolation of nonlinear patterns via manifold learning and dimension reduction, quantification of the circularity of the signal using persistent homology, and comparison of that measure to a bootstrapped null distribution to assess statistical significance (**Figure 1B**). We describe the main steps of the method here; a complete description pertaining to the methods preprocessing, null distribution, and period estimation can be found in the supplement and TimeCycle’s documentation available at https://nesscoder.github.io/TimeCycle/.

**Figure 1:**
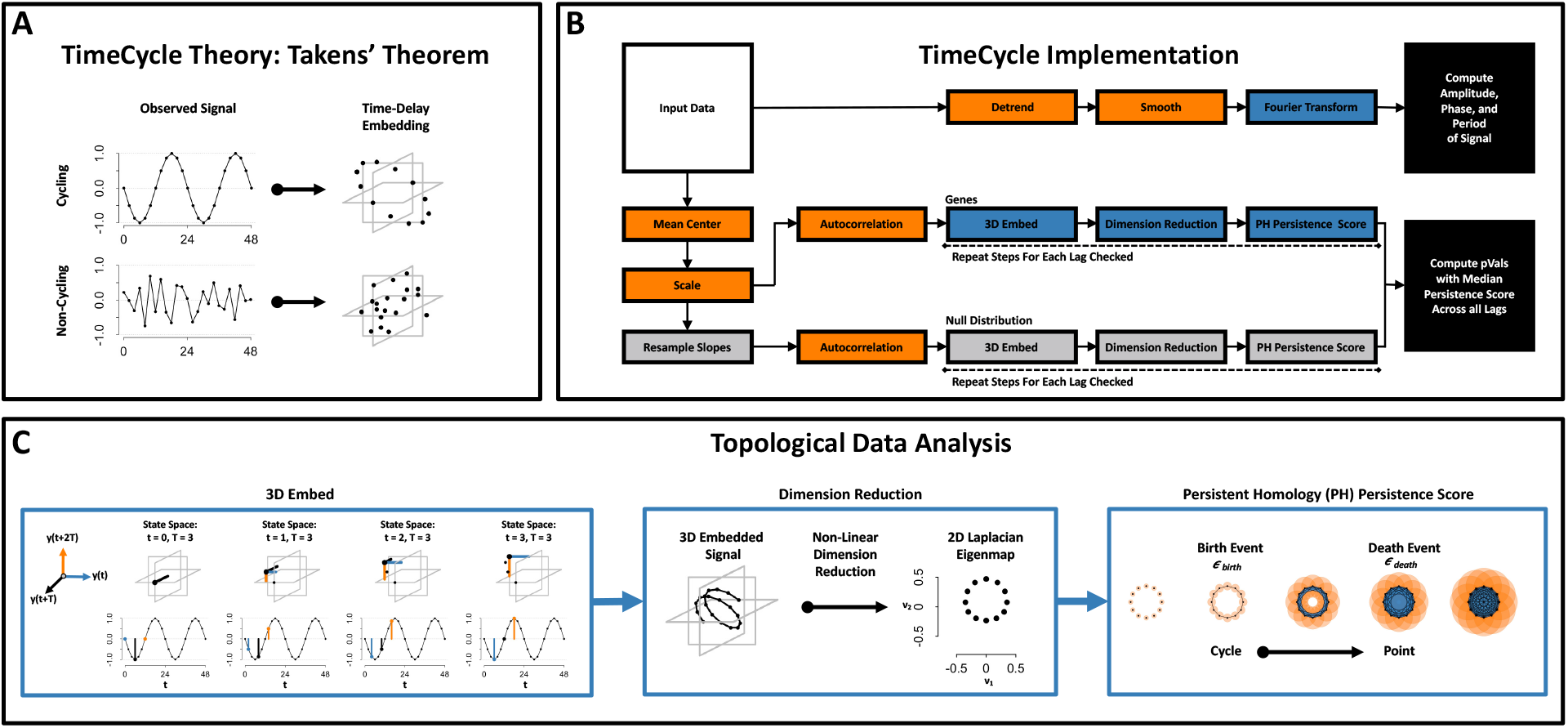
TimeCycle Theory and Implementation. **[A]** Takens’ theorem proves that the dynamics of a periodic signal will exhibit circular patterns in the embedded space. A periodic function (**Top**) embedded in the state space forms a circle. A non-periodic function (**Bottom**) embedded in the state space forms a mass. **[B]** The TimeCycle algorithm assesses statistical significance with respect to a null distribution of persistence scores generated from random time-series with the same marginal distribution of finite differences as the original time-series; this ensures that the random time-series are constrained by the same transcription and degradation rates present in the original data. Period, phase, and amplitude are estimated separately using a Fast Fourier Transform procedure. **[C] Left:** For a given time-series, the method reconstructs the state space using time-delay embedding, a data transformation technique for dynamic systems. **Middle:** Dampened and trending signals form spirals and helices in the embedded Space. Non-linear dimension reduction using Laplacian Eigenmaps yields embedded points preserving the local circular geometry of the 3-D helix in a 2-D space. **Right:** Persistent Homology is used to parameterize the circularity of the 2-D embedding. Points in the embedding are connected if they are at most a distance 2*ϵ* apart, as the radius ϵ increases. A topological cycle appears at some *ϵ*_*birth*_ and disappears at *ϵ*_*death*_, and a persistence score is calculated as Persistence = *ϵ*_death_ − *ϵ*_birth_. Periodic signals have high persistence, while non-periodic signals have low persistence.

### Reconstructing dynamical cycles via Takens’ theorem

TimeCycle exploits Takens’ theorem, a result from dynamical systems theory that proves that the time–delay embedding of a single variable observed over time will reconstruct (up to diffeomorphism) the state space of a multivariate dynamical system [33]. A *d*-dimensional time–delay embedding of a time-series *X* is defined as a representation of that time–series in a *d*-dimensional space where each point is given by the coordinates (*x*_*t*_, *x*_*t*−*τ*_, *x*_*t*−2*τ*_, . . . , *x*_*t*−*dτ*_). As a consequence of Takens’ theorem, a dynamical system with a cycle will exhibit circular patterns in the time–delay space, whereas non-periodic signals will form a mass (**Figure 1A**). Using this logic, TimeCycle reconstructs the state space for each gene using time-delay embedding and quantifies the circularity of the embedding as a measure of the evidence of cycling dynamics.

### Parameter choice and detrending via dimension reduction

Time-delay embedding requires two parameters, the embedding dimensionality *d* and the delay lag *τ*. A perfectly sinusoidal signal will form a circle in *d* = 2 dimensions, and hence a two-dimensional embedding would be sufficient in the ideal case. However, biological signals are often not strictly periodic, but exhibit drifts in the oscillation. A common approach is to detrend the time-series by fitting it to a line and analyzing the residuals [23]. While this removes linear trends, the detrending procedure can introduce false positives [16], as non-rhythmic signals—e.g. sigmoidal and exponential—may appear rhythmic after detrending (**Supplemental Figure 1**). Instead, we observe that any non-periodic component of the signal may be represented by higher dimensions in the embedded space. Hence, we embed the time-series in three dimensions **(Figure 1C: Left)** and use nonlinear dimension reduction to recover the periodic component. An illustrative example is given in **Figure 1C (Middle)**. A drifting oscillation will form a helix in the 3-D embedded space, with the linear trend contributing to the elongation of the helix along one coordinate and the circular component preserved in the other two. TimeCycle uses Laplacian Eigenmaps [34], a nonlinear dimension reduction (NLDR) technique, to project the 3-D embedded data back into a 2-D space, preserving the circular geometry.

This approach has two advantages over linear-fit detrending. First, it is in principle applicable to any type of drift, and is not necessarily confined to removing linear trends. Second, it will only yield a circular pattern in the 2-D space if the periodic component is strong relative to the drift. That is, if the “drift coordinate” (e.g., the axis of the helix in Figure 1C) more faithfully preserves the local geometry of the data than the circular “cycling coordinates”, it will be chosen by the NLDR, and the resulting 2-D representation will not have a detectable circular topology. Together, these features overcome the drawbacks of linear-fit detrending.

The optimal choice for the other parameter, the lag *τ*, is less obvious. While the underlying mathematical theory holds for all *τ* > 0 [33], in practice the presence of noise and short time-series lengths lead to differences in the embedding as a function of *τ* [35]. Because the underlying theory is insensitive to *τ*, it yields no guidance on how to derive an “optimal” lag or set criteria for its selection. A common approach, due to Fraser and Swinney, is to choose *τ* to maximize the independence between embedded points [36]; more recently, Meiss and colleagues proposed a method to choose *τ* that minimizes the curvature [37]. In practice, however, the best choice is often application–dependent. Because our goal here is to detect rhythmicity (rather than fully reconstruct the state space), we sweep through values of *τ*, computing the degree of circularity for each. If a signal is truly rhythmic, on average the state space manifold across lags should be approximately circular.

### Quantification of cycling via persistent homology

We quantify the circularity of the embedded data using *persistent homology* (PH) [38], an algebraic means to measure topological structures (i.e. components, holes, voids, and higher dimensional analogs) in a data set. From each point of the embedded signal, a *d*-sphere of radius *ϵ* is incrementally grown (**Figure 1C: Right**). When a pair of points have intersecting spheres (i.e. are at most 2*ϵ* apart), a line is drawn between those points. When a complete cycle is formed amongst a set of points at some *ϵ*_*b*_, this is defined as a “birth event”: the initial appearance of a topological hole, a closed cycle lying entirely inside the spheres surrounding the embedded data. As *ϵ* is increased, the topological hole eventually closes at a radius *ϵ*_*d*_ defining the “death event”. The persistence score of a topological feature is defined as the difference between the death and birth radii *ϵ*_*d*_ − *ϵ*_*b*_. Several such features, with associated persistence scores, may be present in the data, the largest of which corresponds to the most unambigously periodic dynamics. A rhythmic signal will produce a more circular embedding, resulting in a larger maximum persistence score, than a non-rhythmic signal.

### Significance testing

A common approach for assessing statistical significance in cycling detection algorithms is to compare the statistics of an observed signal to a null model comprising random time series [13, 22, 25]. Such null models may be unrealistic, however, since gene expression in the biological context is constrained by transcription and degradation rates, and an assumption of random, independent time-series may contain changes in gene expression that are biologically unattainable. Instead, TimeCycle’s null model is obtained by permuting the finite differences in gene expression between sampled time-points. The distribution of gene expression changes under the null is thus identical to that of the observed data, ensuring that the resulting random time-series reflect biological transcription and degradation constraints (**Figure 1B**). A persistence score is computed for each resampled time-series to generate a reference distribution of persistence scores under the null (no cycling), conditioned on the observed rates of change. The observed persistence score is then compared to this reference distribution to obtain a *p*-value.

A significant consideration for this method is the computational complexity of the persistence score calculation [*O*(*N* log *N*) in the number of time points for 2-*d* data], which will need to be performed for each gene and each resampled time-series. Indeed, a drawback of the SW1PerS algorithm is its inability into compute a *p*-value due to the computational cost of the method. To overcome this challenge, we devised a scheme to allow for hypothesis testing with improved computational efficiency: prior to the embedding and PH analysis, each gene is mean centered and scaled to unit variance, allowing a common set of permuted time-series to be used for all genes to test the statistical significance of cycling.

### Parameter estimation

Often, researchers desire estimates of the period, phase, and amplitude of the genes detected as cycling. Because the time-delay embedding and PH computation does not provide these estimates, a separate computation in TimeCycle is used to generate these results. To estimate the period, amplitude, and phase of the oscillations (**Figure 1B**), signals are linearly detrended and smoothed via a moving average before fitting the signal to the first three harmonics of the fast Fourier transform (FFT). The period, phase, and amplitude are computed from the FFT fit. (Note that when performing the less computationally costly parameter estimation, the rescaling procedure used in the cycle detection step is omitted to preserve amplitude measures.)

### Replicate time-points

A practical consideration when designing transcriptomic time-series experiment is the trade-off between replicates, sampling resolution, and sampling length in relation to experimental cost [11, 16]. While cycling detection methods do not require replicates [13,22–25,27,30]; technical replicates are necessary for performing additional analysis beyond the scope of cycling detection – e.g. differential expression and differential time-course profiles [39,40]. As such, researchers benefit from algorithms that can incorporate replicates to improve detection of cycling genes [30]. Adhering to best practice guidelines outlined in previous studies [11, 16], TimeCycle averages replicate time-points. The averaged signal is then processed following all steps outlined above.

### Application and evaluation using simulated gene expression data

To comprehensively evaluate TimeCycle’s capabilities across different patterns of temporal gene expression, we applied TimeCycle and four other methods—JTK_CYCLE [22], RAIN [13], GeneCycle [21], and SW1PerS [24]—to synthetic data described in our previous benchmarking work [16]. JTK_CYCLE and RAIN were selected for comparison as the primary methods used in the field of circadian rhythm detection. GeneCycle was selected as an alternative method highlighted in previous benchmarking studies to be robust to outliers [21, 26, 28]. SW1PerS was selected as the only other method utilizing topological data analysis. While other methods such as ARSER [23] and booteJTK [25] were considered based on previous benchmarking studies [16, 26–28], they were ultimately omitted from analysis as they do not natively handle missing data. Synthetic data sets varied in number of replicates (1, 2, 3), sampling intervals (1-h, 2-h, 4-h), sampling length (36-h, 48-h, 72-h, 96-h), and noise levels (10%, 20%, 30%, 40% of signal amplitude), across 11 base waveform shapes. Seven of these 11 shapes were considered cyclic (sine, peak, sawtooth, oscillations about a linear trend, damped, amplified, contractile), and 4 were considered non-cyclic (flat, linear, sigmoid, and exponential). For each waveform in each condition, 1000 “genes” were simulated with varying amplitudes, phases, and shape parameters (e.g., the envelope for damped/amplified waves), yielding in total 11,000 simulated genes for each of the 144 sampling and noise conditions. (Further details of the synthetic data sets can be found in the Methods section and Supplementary Materials of [16].)

### Accuracy and sampling considerations

A summary of TimeCycle results in the synthetic data is found in **Figure 2**. Receiver-operating characteristic (ROC) curves were computed for each of the 144 synthetic data sets, and the area under the curve (AUC) was used to compare classification accuracy across methods. An AUC of 1 represents perfect classification, while an AUC of 0.5 represents a classification no better than pure chance. AUC values are shown as a function of the number of samples collected per time-series. TimeCycle exhibits AUCs ≥ 0.8 across most sampling schemes, with the best performance observed for data sampled every 2-h for 48-h—the scheme primarily used in previous benchmarking studies [13, 16, 24–27].

**Figure 2:**
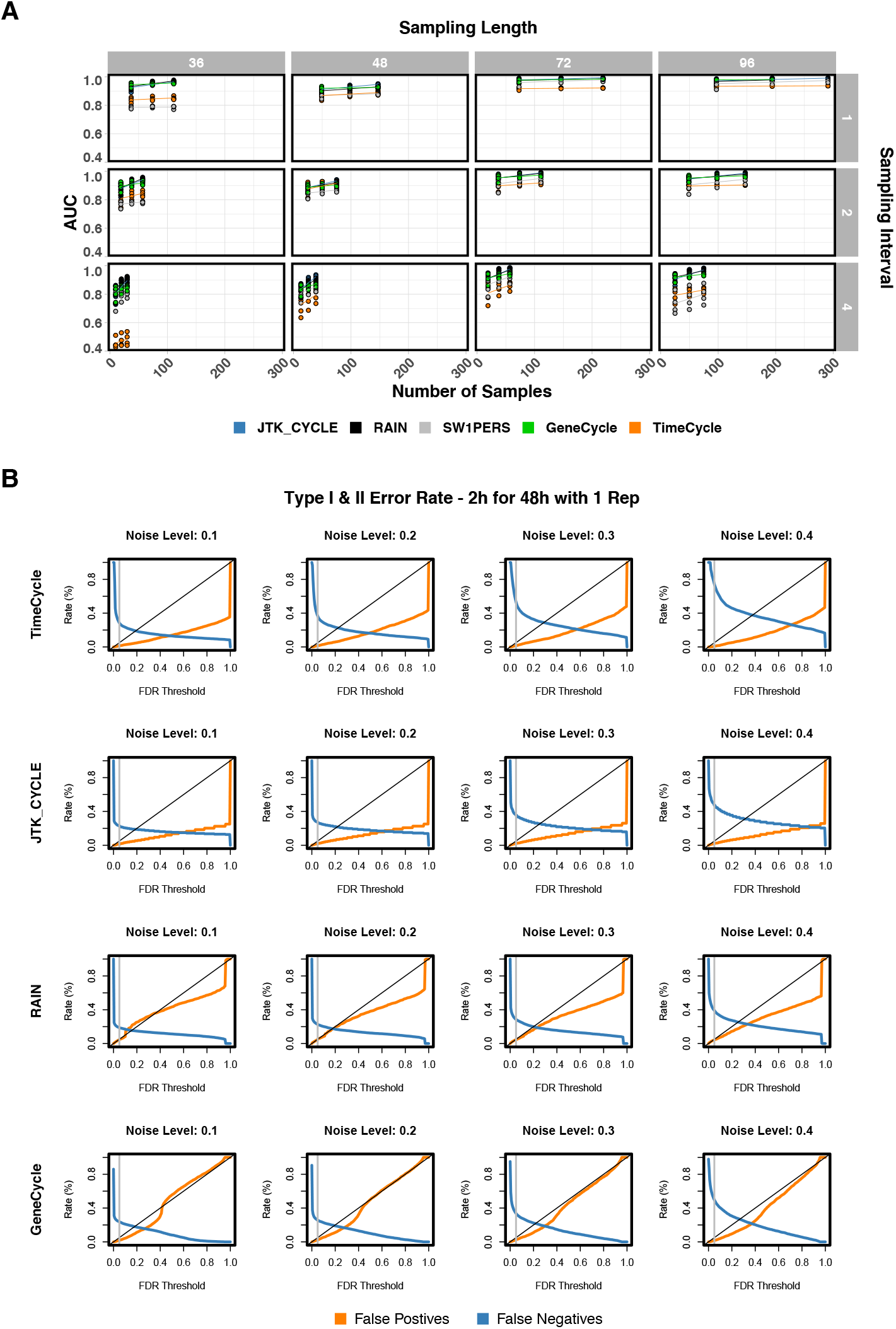
Synthetic Data Overview. **[A]** A total of 144 unique synthetic time-course data sets were generated in R with known ground truth, as described in [16]. Each data set consisted of a different number of replicates (1,2,3), sampling intervals (1-h, 2-h, 4-h), sampling lengths (36-h, 48-h, 72-h, 96-h), and noise levels (10%, 20%, 30%, 40%) as a percentage of the waveform amplitude. AUC scores for each method were computed across all 144 data sets. Lines represent the best linear fit across replicates and noise levels within a specified sampling scheme. Replicate time-series were averaged together for the GeneCycle and SW1PerS algorithm, since neither algorithm has a built-in method for handling replicates. **[B]** Type I and Type II error rates at varying FDR thresholds across methods for synthetic data sampled every 2-h for 48-h with 1 replicate. FDR = 0:05 marked by grey vertical line. (SW1PerS does not compute a p-value, but rather a periodicity score and was thus omitted from analysis.)

Examining the type I and type II error rates relative to the FDR adjusted *p*-values across the varying noise levels, we find that TimeCycle and JTK_Cycle are more conservative in comparison to RAIN and GeneCycle (**Figure 2B**). This can also be seen in **Supplemental Figure 2**, which illustrates that the null distribution of *p*-values is biased toward 1 for both TimeCycle and JTK_Cycle (more strongly biased for JTK_Cycle), whereas small *p*-values are overrepresented under the null for RAIN and GeneCycle (leading to a greater chance of false positives) for non-cycling genes. Across all methods, false negative error rates increase with noise.^*^

Outside the 48-h sampling schemes, TimeCycle exhibits good performance for longer time-series sampled every 1 or 2-hours, but reduced performance for sparser sampling (every 4-h compared to 1-h and 2-h), even for longer time-series. This is attributable to sparsely-sampled manifolds in the reconstructed state space, resulting in an insufficient number of data-points for cycle formation. This suggests that, for a fixed number of samples, denser sampling for a shorter duration may be favorable to longer, sparser sampling.

It is also instructive to examine how the classification accuracy for different waveforms changes with the sampling strategy. We find that for time-series lasting 36h, strong symmetric cyclers are robustly detected, while asymmetric cyclers (such as the sawtooth) do not have sufficient points to close the cycle in the embedded space (that is, observations along the “sharp” rise or fall of an asymmetric waveform may be missed when fewer than two complete periods are assayed, leading to a “C” shape in the embedded space). At 72-h and 96-h, linear trending and damped oscillations are more likely to be classified as non-cycling due to the fact that as the time-series sampling is extended, the trends and dampening become more pronounced, dominating the underlying oscillation. Because these signals are not in fact strictly periodic (returning to precisely the same state at regular intervals), this can be a desirable or undesirable behavior depending on whether the user wishes to classify linear trending and damped oscillations as cycling. Details for an implementation with alternative parameters to detect oscillations with linear trends can be found in TimeCycle’s documentation.

### Comparison to other methods

To compare TimeCycle’s ability to distinguish cycling from non-cycling waveforms to that of other methods, we examined the pairwise comparison of raw *p*-values generated by each method. To simulate results for an ideal data set, comparisons were made using the recommended sampling scheme—every 2-h for 48-h with one replicate—under low (10%) noise conditions (**Figure 3**). (Similar plots for varying noise levels can be found in **Supplemental Figures 3–5**).

**Figure 3:**
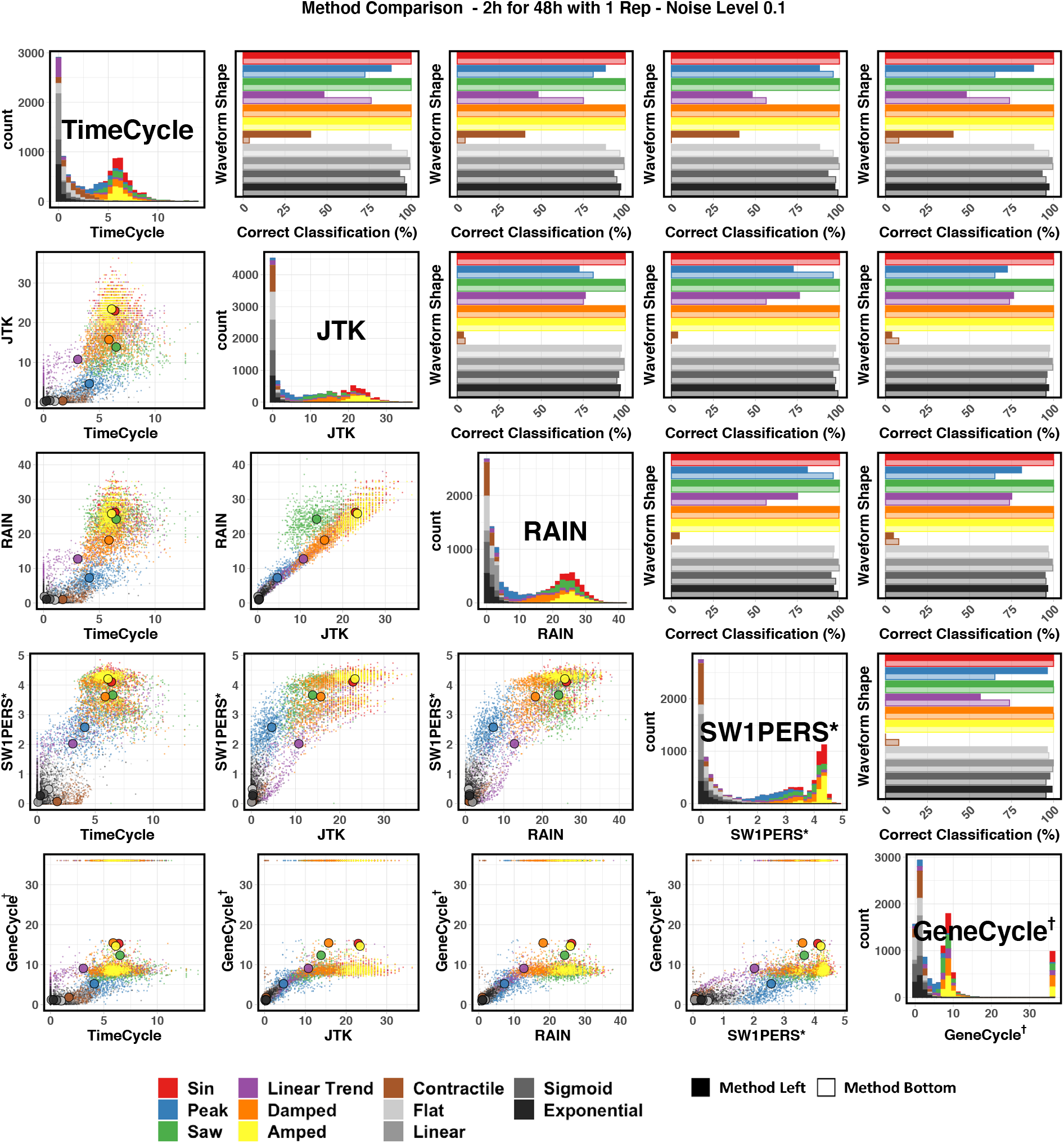
48 Hour Synthetic Data Method Comparison. Comparison of TimeCycle, JTK_CYCLE, RAIN, SW1PerS, and GeneCycle results for synthetic data of various waveform shapes. **LOWER TRIANGLE:** scatterplot of −log_10_ *p* for each synthetic gene. The larger colored points represent the average −log_10_ *p* designated by waveform shape. **DIAGONAL:** Histogram of −log_10_ *p* p for each synthetic gene by method. **UPPER TRIANGLE:** Classification accuracy comparison by waveform category. Darker bars correspond to the method listed in the row; lighter bars correspond to the method listed in the column. Each value corresponds to the fraction of correctly classified genes (true positive for cycling/non-cycling as appropriate) for a given waveform type, using a classification threshold obtained from Younden’s *J* index for the ROC across all genes. * SW1PerS does not compute a *p*-value, but rather a periodicity score. † GeneCycle results of *p* = 0 were set to machine precision (2:2 · 10^−16^) for visualization purposes.

The lower triangle of **Figure 3** depicts the pairwise scatterplot of the − log_10_ *p* for each synthetic gene across methods, with the larger points designating the average − log_10_ *p* for a given waveform shape. (Note that, as previously discussed, SW1PerS does not compute a *p*-value; instead, the periodicity score is shown.) The results are generally well-correlated, with non-cycling waveforms clustering at low significance and sinusoidal waveforms ranked high; differences between methods are most noticeable for trending, sawtooth, and contractile waveforms. Given the algorithmic similarities of RAIN/JTK_CYCLE (via the Jonckheere-Terpstra test) and TimeCycle/SW1PerS (via topological analysis), it is unsurprising that these methods are more highly correlated within method pairings than across methodological approaches.

The upper triangle of **Figure 3** measures the fraction of correctly classified signals for each waveform shape, ie, the conditional probability of correctly classifying the gene as cycling or non-cycling given the waveform. Here, the decision threshold for classifying a given gene as cycling was chosen via Youden’s *J* statistic [41] computed from AUC over all genes. For the sampling scheme of 48h every 2h, TimeCycle’s accuracy was comparable to other methods for most waveform shapes with three exceptions: First, TimeCycle shows improved performance in detecting peaked and contractile waveforms across methods when samples are taken every 2h across 48h. Second, TimeCycle also shows improved detection of peaked waveforms in comparison to JTK_CYCLE, RAIN, and GeneCycle, with a slight decrease in comparison to SW1PerS. Third, TimeCycle shows decreased performance in detecting linear trends across all methods. To mitigate this shortcoming, alternative parameters for the improved detection of genes with linear trends can be found in TimeCycle’s documentation. While TimeCycle outperforms other methods in certain circumstances, our result suggest that no method is uniformly best at picking up all types of circadian dynamics under all sampling schemes.

### Missing data and outliers

Since missing data often occur as a result of sequencing errors with greater probability as the sample size increases [32], researchers benefit from algorithms that can handle missingness. TimeCycle imputes the missing points via the linear interpolation algorithm as described by Moritz *et al*. in their imputeTS R package [42]. However, by introducing imputed data, one also introduces new assumptions—specifically, smoothness in the trajectory and hence small gene expression changes before/after the missing observation. These properties must be shared by the null model to ensure that the significance testing is not biased by the imputation. Hence, we perform the previously–described resampling procedure post-imputation, effectively taking the imputation assumptions into account when defining the null distribution.

To investigate how TimeCycle performs when data are missing, we simulated missingness in the synthetic data sets (**Figure 4**). For each of the 11,000 genes, we randomly removed points sweeping from 0% missingness until only 2 points remained. The AUC was computed at each missingness value (**Figure 4A**). At 50% missingness in the synthetic data, TimeCycle shows robust classification with a slow decrease in AUC for each noise level. We observe that the difference in AUC between noise levels is greater than the degradation in AUC as a function of missingness (**Figure 4A & B**). The differences are primarily attributable to decreases in sensitivity, with high levels of noise and missingness compromising the ability to detect cycling genes. We find little decrease in specificity, confirming that the “smoothing” induced by the imputation procedure does not generate false positives. Missing data analyses for JTK_CYCLE, RAIN, and GeneCycle can be found in **Supplemental Figures 6–8**.

**Figure 4:**
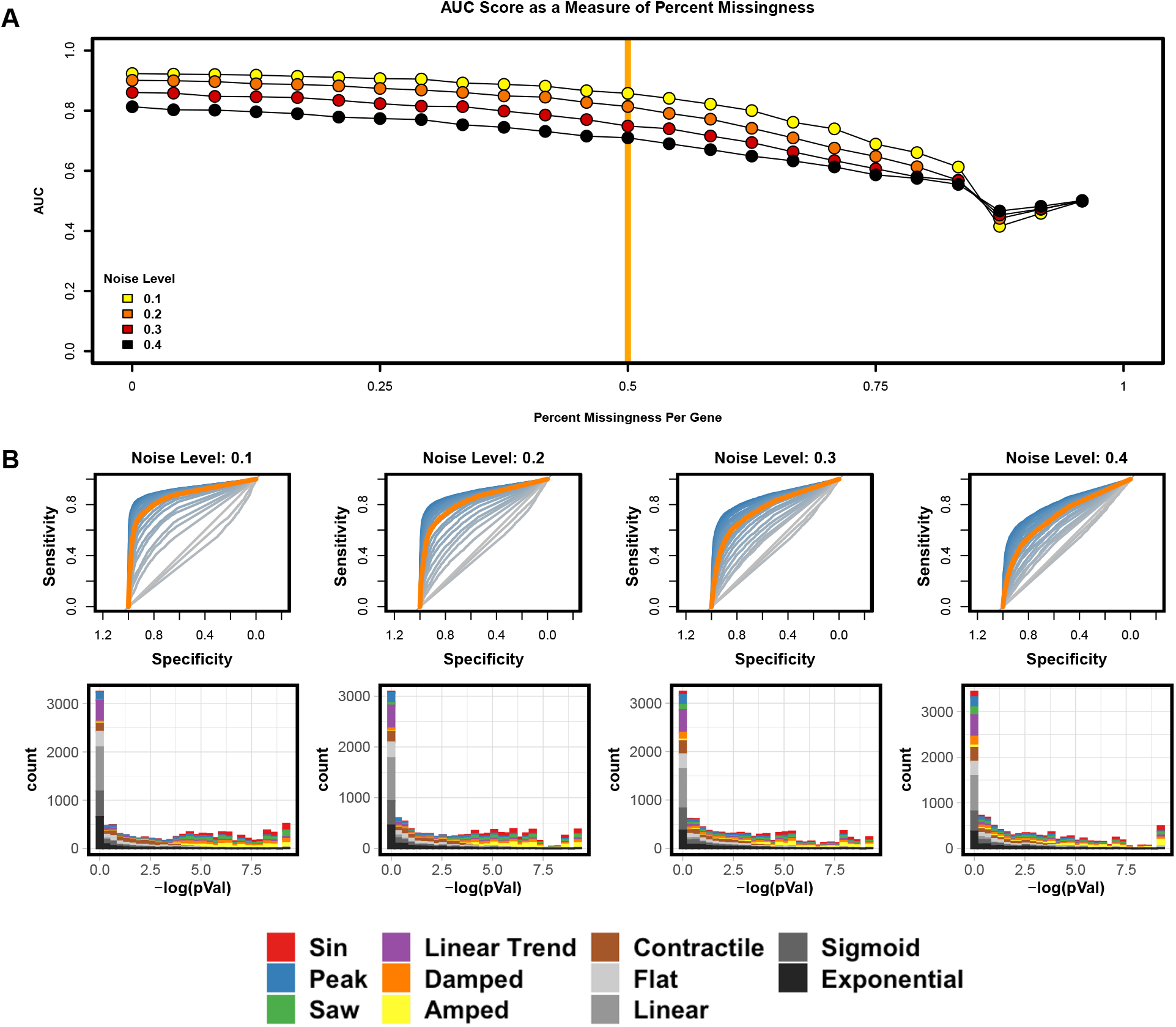
Missing Data Results. **[A]** TimeCycle AUC scores across varying levels of percent missingness per gene when sampled every 2-h for 48 h with one replicate. Results are shown for each level of noise. 50% missingness is highlighted by the vertical orange line. **[B] TOP:** ROC curves for each percent missingness depicted in the AUC score plot in panel A above. ROC curves scaled from Blue (0% missingness) to grey (96% missingness). 50% missingness is again highlighted by the orange curve. **BOTTOM:** Histogram of −log(*p*) at the different noise levels corresponding to the orange 50% missingness ROC plot.

As an additional check of method robustness, we performed an outlier analysis across all 36 variants of the 48-h sampling lengths. Outliers were injected into the time–series at a rate of 1% of all sampled time–points and were drawn from a uniform distribution between [*μ* − 4*σ*, *μ* − 3*σ*] and [*μ* + 3*σ*, *μ* + 4*σ*] of the diurnal mean (*μ*) for each time–series, following previous studies [22, 43]. TimeCycle exhibited a very slight and non-significant decrease in average AUC±sd across all noise levels (0.86±0.05 vs 0.87±0.05, with and without outliers, respectively); other methods also remained unchanged (**Supplemental Figure 9**). As expected from the unchanged AUC, the type I and type II error rates were also unaltered. This suggests that the performance of TimeCycle, as well as other methods, is robust to outliers.

### Application to biological data: reproducibility analysis

While synthetic data has the advantage of known ground truth, it also has the drawback of not necessarily being representative of the real biological data sets. On the other hand, measuring a method’s accuracy using real data is limited, as the ground truth is not generally known. Instead, one may test the reproducibility of the results, under the assumption that a true biological signal should be consistently detected across multiple studies of the same condition.

To this end, we took a “cross-study concordance” approach in which we tested the ability of each method to consistently characterize a set of 12,868 genes measured in three independent mouse liver time-series expression sets [1, 44, 45]. We expect that a method that accurately detects cycling genes should do so reproducibly in all three data sets, whereas a method that is no better than chance will yield divergent results in the three data sets. Moreover, the genes detected as cycling in common across the data set should exhibit similar dynamics across all three data sets. By contrast, if a method is overly permissive, leading to a high false-positive rate, genes may be classified as “cycling” across data sets without actually having any commonality in their dynamics. We evaluate both of these by (i) examining the overlap in genes classified as cycling in the various data sets (following the Methods of [16]) and (ii) computing the rank correlation *ρ* of the expression profiles of the genes detected as cycling at FDR < 0.05 and LogFC > 2. Together, our analysis quantifies whether the cycling detection is reproducible.

The data sets used in this analysis were comparable, but independent time-series studies of gene expression in mouse liver, referred to hereafter as Hogenesch [44], Hughes [45] and Zhang [1]. These data sets are summarized in **Table 1**. The Hogenesch data set [44] was downsampled to two data sets sampled every 2-h for 48-h (Hogenesch 2A and Hogenesch 2B) to be consistent with the sampling schemes of the Hughes 2012 [45] and Zhang 2014 [1] studies. Following application of TimeCycle, genes with FDR < 0.05 were classified as cycling; the overlap in the cycling genes for the various data sets is given (**Figure 5A: Top**). Comparing the distribution of the FDR-corrected *p*-values for the known circadian genes relative to all genes shows a statistically significant shift across all four datasets, with circadian genes more likely to have significant *p*-values (**Figure 5A: Bottom**), confirming the intuition that known circadian genes should show enriched cycling. Furthermore, the known circadian genes Per1, Cry1, Npas2, and Clock were detected across multiple studies.

**Table 1:** Experimental Design Biological Data. Three mouse liver time-series expression sets (Hogenesch 2009, Hughes 2012, Zhang 2014) where analyzed from the Gene Expression Omnibus database (GEO). In each experiment, Wildtype C57BL/6J mice where entrained to a 12 hour light, 12 hour dark environment for a week before being released into constant darkness, a standard practice in the field for assessing circadian function. The Hogenesch study sampled mice every 1 hour for 48 hours, while the Hughes and Zhang Studies sampled every 2 hours for 48 hours. Other variations in the data sets include microarray platform, ZT time of sampling, and number of mice sampled per time point. Adapted from [16].

|  | **Hogenesch 2009** | **Hughes 2012** | **Zhang 2014** |
| --- | --- | --- | --- |
| **Genotype** | Wildtype C57BL/6J | Wildtype C57BL/6J | Wildtype C57BL/6J |
| **Age** | 6-weeks | Not Specified | 6-weeks |
| **Entertainment Condition** | 12h : 12hLD → DD | 12h : 12hLD → DD | 12h : 12hLD → DD |
| **Entrainment Length** | 1 Week | 1 Week | 1 Week |
| **Microarray Platform** | Affy Mouse 430 2.0 | Affy MoEx 1.0 ST | Affy MoGe 1.0 ST |
| **Sampling Scheme** | Every 1-h for 48-h | Every 2-h for 48-h | Every 2-h for 48-h |
| **ZT Start Time** | ZT 18 | ZT 0 | ZT 18 |
| **Samples Per Time Point** | 3-5 Male | 2 Male and 2 Female | 3 Male |
| **GEO Accession #** | GSE11923 | GSE30411 | GSE54650 |

**Figure 5:**
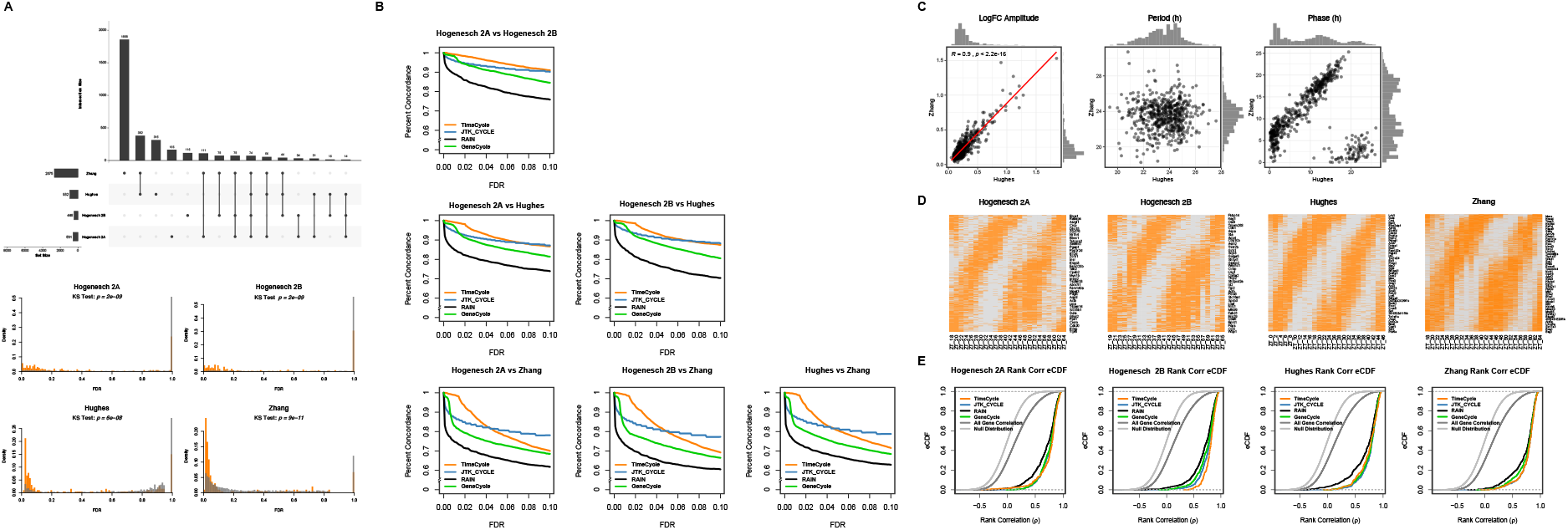
TimeCycle Biological Data Results. **[A] TOP:** Upset plot showing overlap between significant cycling genes sets detected by TimeCycle at an FDR < 0:05 across 3 distinct mouse liver time-series data sets as described in [16].The Zhang and Hughes data sets were sampled every 2-h for 48-h. The Hogenesch study, sampled every 1-h for 48-h, was downsampled into two data sets sampled every 2-h for 48-h (Hogenesch 2A and Hogenesch 2B). **BOTTOM:** Distribution of FDR-adjusted *p*-values of the known circadian genes (orange) versus all genes (grey). Circadian genes were extracted from the CGDB database [48]. Only circadian genes that were experimentally validated in mouse liver tissue through low-throughput methods were included in the analysis. **[B]** Percentage of genes concordantly called cycling or non-cycling across studies at varying FDR thresholds in each pair of studies; comparisons to other methods are also shown. **[C]** LogFC amplitude, period, and phase scatterplot comparison of genes identified as cycling in both the Zhang and Hughes data sets with an FDR < 0:05. **[D]** Heatmap of the cycling genes detected in each data set with an FDR < 0:05 ordered by phase. Orange and grey represents gene expression above and below diurnal mean, respectively. **[E]** Rank correlation CDFs of genes identified by each method with an FDR < 0:05 and LogFC < 2 in one data set compared to their pairwise expression in the other data sets. The null distribution represents the rank correlation of a random sampling of all 12,868 genes with replacement in the pairwise comparisons of all data sets. All gene correlation represents the rank correlation on a per gene basis for all 12,868 genes in the pairwise comparisons of all data sets.

To investigate whether the reproducibility was robust to the (somewhat arbitrary) choice of significance threshold, we calculated the percent of genes concordantly classified as cycling or noncycling for each pair of studies as a function of the FDR significance threshold (**Figure 5B**). The concordance is generally higher for TimeCycle relative to other methods at conservative significance thresholds ≤ 0.05. This implies that genes identified as cycling by TimeCycle are likely to be reproducible.

The Hughes and Zhang studies differed in the sampling start phase by 6-h, presenting an opportunity to examine whether genes detected as cycling in both had similar estimated periods and amplitudes, but differing phases, as would be expected if the cycling detection is accurate [46]. We find that genes detected as cycling in both the Zhang and Hughes data sets with an FDR< 0.05 also had highly reproducible amplitudes across the various data sets (Pearson correlation *r* = 0.9, **Figure 5C**). We also find that genes identified as cycling clustered around a period of 24 hours, indicating that these are indeed circadian oscillators. Finally, plotting the computed phase of genes detected reflects the 6-h phase shift (modulo 24h) as expected (**Figure 5C**). Heatmaps of cycling genes detected across data sets are shown (**Figure 5D**).

We then examined more broadly whether genes that are detected as cycling across different studies exhibit similar dynamics, and compared TimeCycle’s results to other methods. A method that has a high rate of false-positives may indeed identify many genes as cycling in common across studies, but by chance rather than detecting a meaningful signal. Hence, we assess whether the genes that are identified as cycling in multiple studies of the same tissue also have reproducible dynamics. For each study and each gene, we computed the rank correlation of its time-series in the reference study with the same gene’s time-series in the other studies. We expect that genes under true circadian control should exhibit more correlated dynamics amongst the studies than the set of all genes. We thus computed empirical CDFs of the correlations for genes identified as cycling (FDR < 0.05) by each method^†^ (**Figure 5E**). In this figure, the null distribution represents the rank correlation of a random sampling of all 12,868 genes with replacement in the pairwise comparisons of all data sets, and the all-gene correlation represents the rank correlations for all 12,868 genes in the pairwise comparisons of all data sets. TimeCycle, JTK_CYCLE, and GeneCycle show a statistically significant shift in the distribution of rank correlations in comparison to RAIN, the all-gene correlation, and the null distribution in all data sets (all *p* ≤ 2.2 · 10^−16^, Kolmogorov-Smirnov test). We conclude from this that TimeCycle reliably detects cycling dynamics across multiple studies of the same tissue.

## Discussion

We have presented TimeCycle, a new method that leverages results from dynamical systems theory and topology to detect patterns of cyclic expression in time-series experiments. TimeCycle reconstructs the state space for the dynamical system governing each gene using time-delay embedding, and quantifies how cyclic the embedding is using persistence homology. Statistical significance is assessed by comparing the persistence scores to those that obtain from a resampled null model. TimeCycle accurately detects rhythmic transcripts in both synthetic and real biological data, and is robust to missingness, noise, and non-cyclic components in the dynamics.

A few methodological innovations distinguish TimeCycle from other cycling-detection algorithms. In contrast to methods that compare gene expression profiles to templates of expected cycling patterns, TimeCycle reconstructs the underlying dynamical system directly from the observed data. This enables TimeCycle to articulate more complex dynamics than can be easily considered using template-based approaches. Additionally, the method to construct the null distribution is both computationally efficient and biologically representative. SW1PerS (a prior method that also used persistence homology rather than a template) did not implement hypothesis testing due to the computational cost, while template-based methods that implement hypothesis testing do so by resampling the time-series in a manner that can generate biologically implausible null models. TimeCycle is thus an improvement in both regards.

From a practical standpoint, we identified strengths and weaknesses of TimeCycle’s ability to detect cycling transcripts under varying conditions. We find that TimeCycle is better able to detect sharply peaked waveforms and waveforms where the period appears variable, while JTK_Cycle, RAIN, and GeneCycle are more robust with respect to linear trends. This is in keeping with prior work [16] demonstrating that no cycling detection method is consistently “best” for all genes. Instead, it is incumbent upon the researcher to consider the patterns of gene expression that are of the greatest interest and choose a method accordingly.

The results also highlight the importance of constructing biologically representative null models. By resampling the finite differences from the gene expression time-series to construct a null distribution of persistence scores, TimeCycle tests whether an observed gene has a stronger cycling behavior than expected by chance, conditioned upon the speed at which the expression is capable of changing. This improves upon SW1PerS, which does not compute a *p*-value, and also upon methods such as RAIN that randomize the time-series itself. We note that this method for constructing the null distributions could also be adapted for other methods, and emphasize the need for method developers to consider biological constraints when devising null models.

A practical consequence of TimeCycle’s methodological features is that the genes detected as cycling by TimeCycle are highly reproducible. Applied to three independent studies of mouse live gene expression, TimeCycle consistently identified genes as cycling in multiple studies, and those genes were shown to exhibit reproducible dynamics.

Finally, we note that the experimental sampling design remains a crucial factor for the reliability of any cycling detection method. As with other methods [11, 16], TimeCycle performs best when applied to time-series spanning 48-h sampled every 2-h, and is considerably less accurate for shorter and sparser sampling. These findings underscore the important role that experimental design — not only the method choice — plays in the analysis of circadian data.

## Methods

### Generating synthetic data

A total of 144 unique synthetic time-course data sets, each comprising 11,000 expression profiles, were generated in R using the code outlined in [16]. Each data set consisted of a different number of replicates (1, 2, 3), sampling intervals (1-h, 2-h, 4-h), sampling lengths (36-h, 48-h, 72-h, 96-h), and noise levels as a percentage of the wave form amplitude (10%, 20%, 30%, 40%). Within each condition, 11 base waveforms were simulated to mimic expression patterns observed in nature: periodic patterns, nonperiodic patterns, and dynamics that have a cyclic component but do not meet the strict definition of periodicity. Seven of these 11 shapes were considered cyclic (sine, peak, sawtooth, linear trend, damped, amplified, contractile), and 4 were considered noncyclic (flat, linear, sigmoid, and exponential). Further details may be found in [16].

### Processing synthetic data

All 144 synthetic data sets were processed by all four cycling detection methods (TimeCycle, JTK_CYCLE [22], RAIN [13], GeneCycle [21], and SW1PerS [24], using each method’s recommended parameter settings as defined by the method’s documentation. Since GeneCycle and SW1PerS do not have a built-in function for dealing with replicates, replicates were averaged together, following the recommended common practice in the field [11, 16]. TimeCycle, JTK_CYCLE, and RAIN used the replicate procedures recommended in their documentation. See the Supplement and https://github.com/nesscoder/TimeCycle-data for a complete list of the experimental parameters and associated source code.

### Computing ROC, AUC, and percent correct classification

For each method across all 144 synthetic data sets and missing data analysis, the receiver operating characteristic (ROC) and accompanying area under the curve (AUC) were computed using the pROC R package [47]. The optimal threshold for computing each methods’ percent correct classification is defined by the Youden’s *J* statistic [41], again computed using pROC. For each waveform shape at the classification threshold defined by the Youden’s *J* index, the percent correct classification was computed by dividing the number of synthetic genes called as (non-)cyclers out of the total possible (non-)cyclers.

### Missing data analysis

To evaluate TimeCycle’s ability to handle missing data, we analyzed synthetic data sampled every 2-h for 48-h with one replicate at varying noise levels with a range of missing values. For each of the 11,000 genes in each data set, time-points were randomly removed by sweeping from 0% missingness until only 2 points remained. ROC plots and AUC scores depicted in **Figure 4** were computed as described above.

### Outlier analysis

To evaluate TimeCycle’s ability to handle outliers, we analyzed 36 synthetic datasets all with a 48-h sampling length. Each data set consisted of a different number of replicates (1, 2, 3) and sampling intervals (1-h, 2-h, 4-h). Following [22] and [43], for all sampled time–points, outliers were injected into the time–series at a rate of 1%. Outliers were drawn from a uniformly distribution between [*μ* − 4*σ*, *μ* − 3*σ*] and [*μ* + 3*σ*, *μ* + 4*σ*] of the diurnal mean (*μ*) for each time–series [43].

### Preprocessing and analysis of microarray data

Microarray data was preprocessed as described in *Methods—Preprocessing Microarray Data* of [16]. (See https://github.com/nesscoder/TimeTrial for associated source code.)

To characterize the effects of sampling schemes using real data, the three data sets were processed to ensure comparability across data sets. The Hughes 2012 and Zhang 2014 data sets were sampled every 2-h for 48-h. The Hogenesch 2009 data set, comprising data sampled every 1-h for 48-h, was down-sampled into 2 data sets sampled ever 2-h for 48-h. All four data sets (2 original and 2 down-sampled) were processed by TimeCycle, JTK_CYCLE, RAIN and GeneCycle, using each method’s recommended parameter settings as defined by the method’s documentation.^‡^ Further details may be found in the Supplement.

### Gene dynamics reproducibility assessment via rank correlation CDF

The distribution of Spearman rank correlations were computed by comparing genes identified by each method with an FDR < 0.05 and LogFC < 2 in one data set (e.g. Hogenesch 2009, Hughes 2012, Zhang 2014) with their pairwise expression in the other data sets. Before computing the correlation — to account for the 6-h phase shift in sampling start time between the Hughes dataset (start 0_*ZT*_) and the Hogenesch/Zhang (start 18_*ZT*_) data sets — the last three time points (e.g. 6-h) in the Hughes data set were moved to the start of the time-series (**Figure 5D**). The distribution of cycling gene correlations was compared to both the all-gene correlations and a null distribution of correlations. The all-gene correlation represents the rank correlation on a per gene basis for all 12,868 genes, whether or not they were identified as cycling. The null distribution of correlation coefficients was constructed by randomly resampling the 12,868 genes in each data set (thereby generating correlation coefficients from different, rather than the same, genes).

## Acknowledgements

Research reported in this publication was supported by the NSF-Simons Center for Quantitative Biology at Northwestern University, an NSF-Simons MathBioSys Research Center. This work was supported by a grant from the Simons Foundation/SFARI (597491-RWC) and the National Science Foundation (1764421). The content is solely the responsibility of the authors and does not necessarily represent the official views of the National Science Foundation and Simons Foundation.

This research was supported in part through the computational resources and staff contributions provided for the Quest high performance computing facility at Northwestern University which is jointly supported by the Office of the Provost, the Office for Research, and Northwestern University Information Technology.

## Author contributions

E.N.C. and R.B. devised the TimeCycle method and designed the study; E.N.C developed the TimeCycle package, analyzed the data, and produced the tutorials/documentation; E.N.C. and R.B. wrote the paper.

## Availability of source code and data

### TimeCycle Package

A fully documented open-source R package implementing TimeCycle is available at: https://nesscoder.github.io/TimeCycle/.

### Data and analysis used in this manuscript

Data and source code to reproduce the analysis and figures in this paper are available at: https://github.com/nesscoder/TimeCycle-data.

## Footnotes

* SW1PeRS was omitted from the type I and type II error analysis as it does not produce *p*-values, but rather a periodicity score.

† Since SW1PerS does not produce a *p*-value to classify genes as cycling/non-cycling, it was omitted from this comparison.

‡ SW1PerS was omitted from the microarray analysis since the algorithm does not produce a *p*-value and thus could not be meaningfully compared to the other methods.

