## Supplemental Figures for "TimeCycle: Topology Inspired MEthod for the Detection of Cycling Transcripts in Circadian Time-Series Data"

May 5, 2021

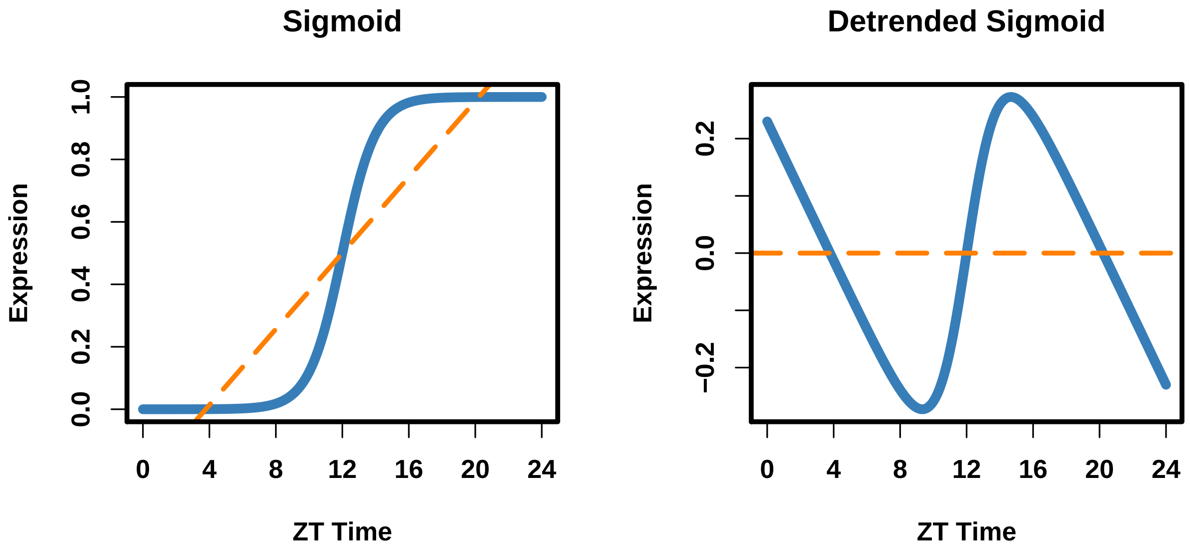


**Figure S1: Linear Detrending of Non-Rhythmic Signals May Induce Rhythmicity | [LEFT]** Sigmoidal waveform (blue line) with a linear fit (orange line). **[RIGHT]** Sigmoidal waveform (blue line) with the linear trend removed (orange line).

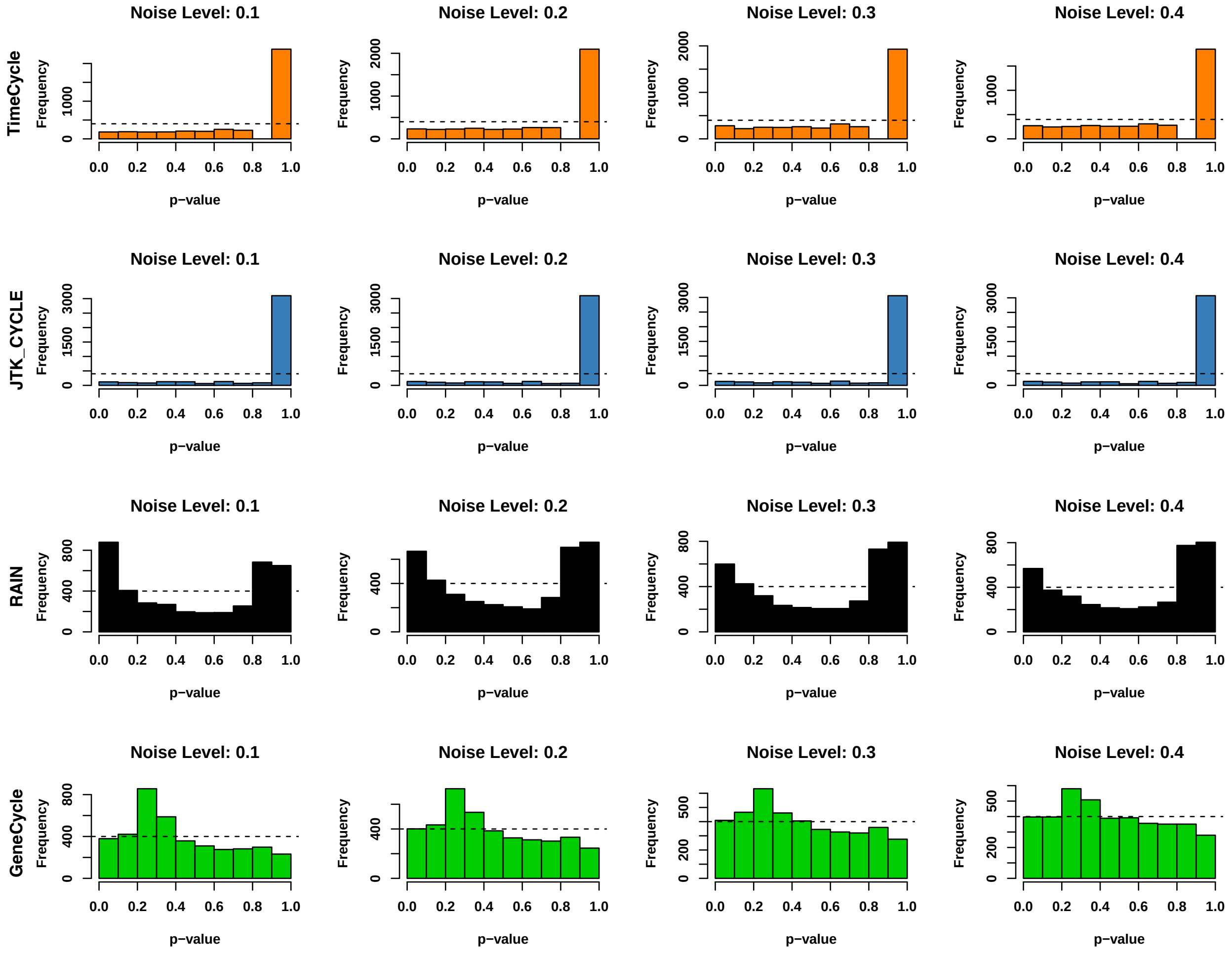


**Figure S2:** ***p*-value Distribution for non-cycling Waveforms** | *p*-value distributions of non-cycling waveforms across methods and noise levels (10%, 20%, 30%, 40%) as a percentage of the waveform amplitude. Gap at 0.9 in TimeCycle's *p*-value consequence of finite bootstrapped sampling. Dashed line represents a uniform distribution of *p*-values under the null.

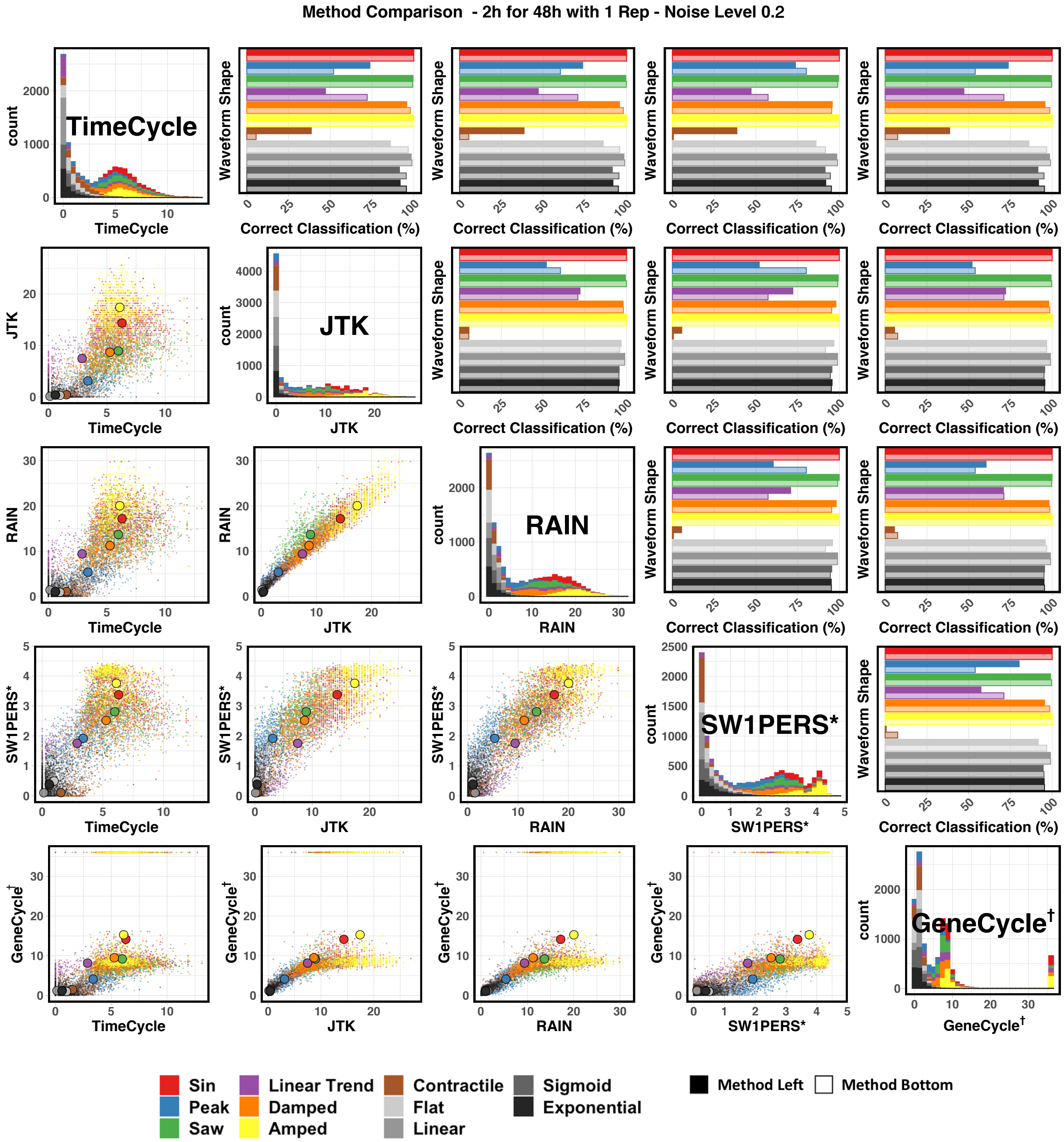


**Figure S3: 48 Hour Synthetic Data Method Comparison - Noise Level 0.2** | Comparison of TimeCycle, JTK_CYCLE, RAIN, SW1PerS, and GeneCycle results for synthetic data of various waveform shapes. **LOWER TRIANGLE:** scatterplot of $-\log_{10} p$ for each synthetic gene. The larger colored points represent the average $-\log_{10} p$ designated by waveform shape. **DIAGONAL:** Histogram of $-\log_{10} p$ for each synthetic gene by method. **UPPER TRIANGLE:** Classification accuracy comparison by waveform category. Darker bars correspond to the method listed in the row; lighter bars correspond to the method listed in the column. As AUC scores are not well defined for subsetted data, $p$-value threshold cutoffs for computing the percent correct classification were based on an optimal ROC threshold as computed by the Younden's $J$ Index. *SW1PerS does not compute a $p$-value, but rather a periodicity score. †GeneCycle results with $p = 0$ were set to machine precision ($2.2 \cdot 10^{-16}$) for visualization.

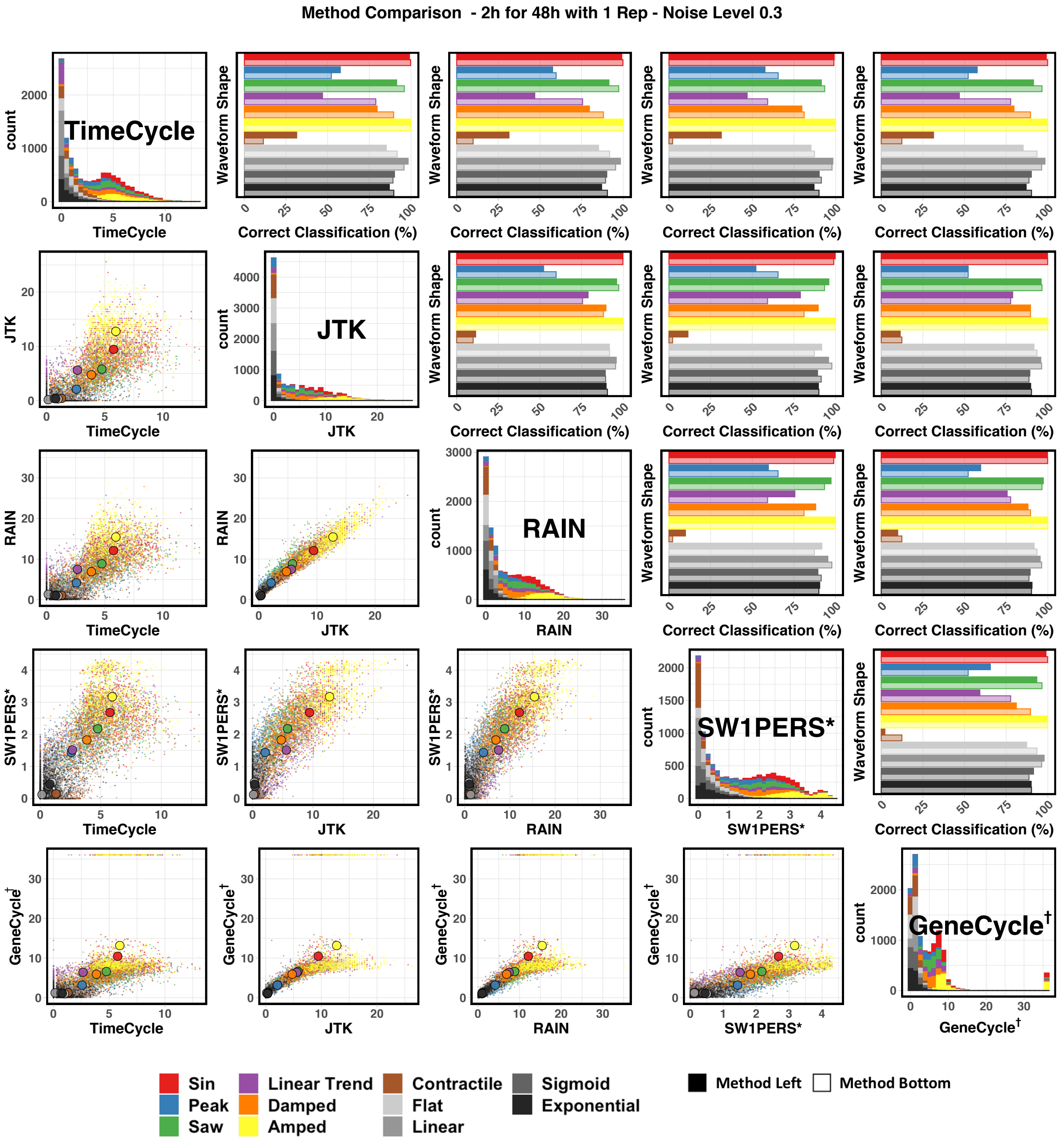


**Figure S4: 48 Hour Synthetic Data Method Comparison - Noise Level 0.3** | Comparison of TimeCycle, JTK_CYCLE, RAIN, SW1PerS, and GeneCycle results for synthetic data of various waveform shapes. **LOWER TRIANGLE:** scatterplot of $-\log_{10} p$ for each synthetic gene. The larger colored points represent the average $-\log_{10} p$ designated by waveform shape. **DIAGONAL:** Histogram of $-\log_{10} p$ for each synthetic gene by method. **UPPER TRIANGLE:** Classification accuracy comparison by waveform category. Darker bars correspond to the method listed in the row; lighter bars correspond to the method listed in the column. As AUC scores are not well defined for subsetted data, $p$-value threshold cutoffs for computing the percent correct classification were based on an optimal ROC threshold as computed by the Younden's $J$ Index. *SW1PerS does not compute a $p$-value, but rather a periodicity score. †GeneCycle results with $p = 0$ were set to machine precision ($2.2 \cdot 10^{-16}$) for visualization.

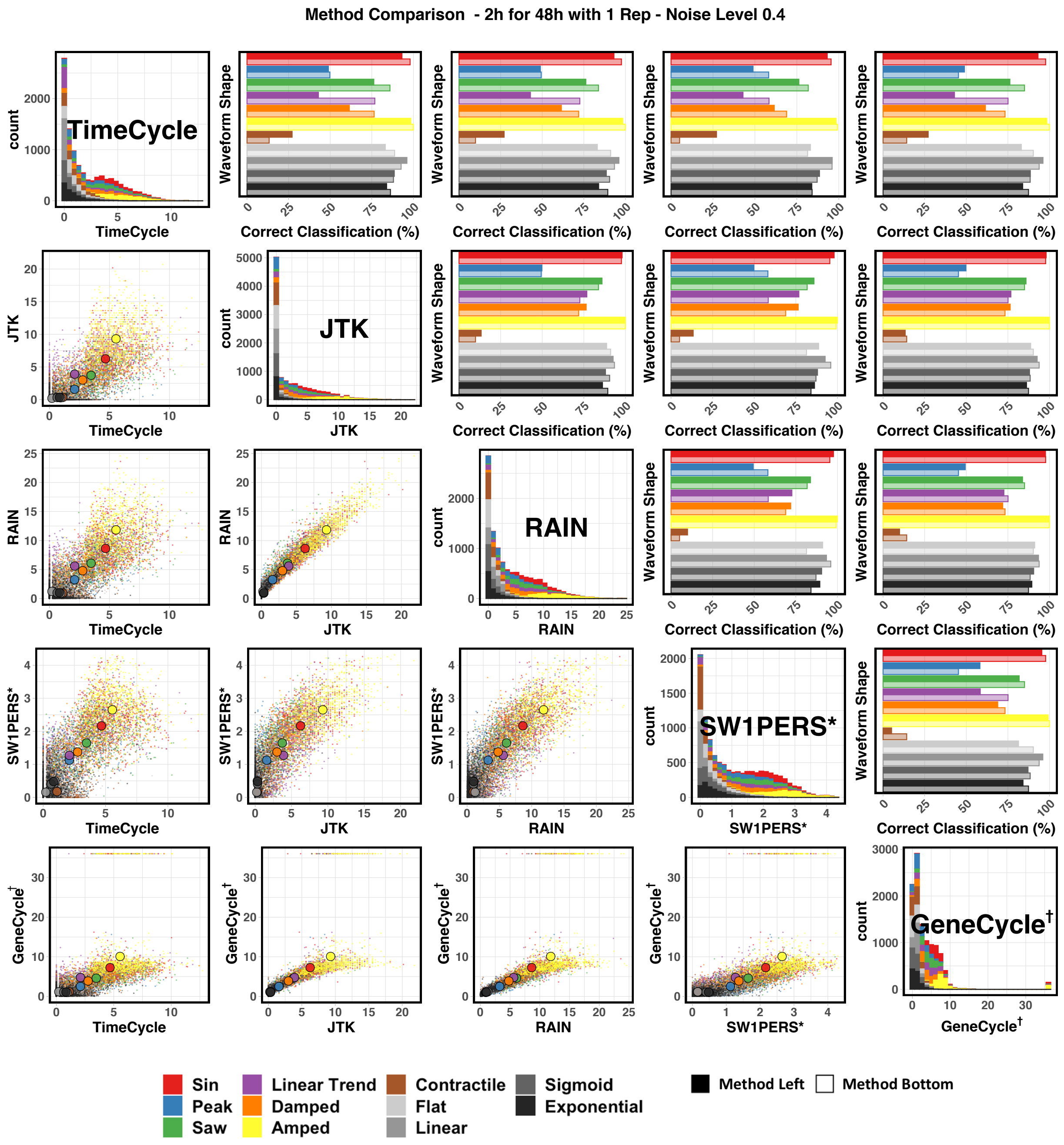


**Figure S5: 48 Hour Synthetic Data Method Comparison - Noise Level 0.4** | Comparison of TimeCycle, JTK_CYCLE, RAIN, SW1PerS, and GeneCycle results for synthetic data of various waveform shapes. **LOWER TRIANGLE:** scatterplot of $-\log_{10} p$ for each synthetic gene. The larger colored points represent the average $-\log_{10} p$ designated by waveform shape. **DIAGONAL:** Histogram of $-\log_{10} p$ for each synthetic gene by method. **UPPER TRIANGLE:** Classification accuracy comparison by waveform category. Darker bars correspond to the method listed in the row; lighter bars correspond to the method listed in the column. As AUC scores are not well defined for subsetted data, $p$-value threshold cutoffs for computing the percent correct classification were based on an optimal ROC threshold as computed by the Younden's $J$ Index. *SW1PerS does not compute a $p$-value, but rather a periodicity score. †GeneCycle results with $p = 0$ were set to machine precision $(2.2 \cdot 10^{-16})$ for visualization.

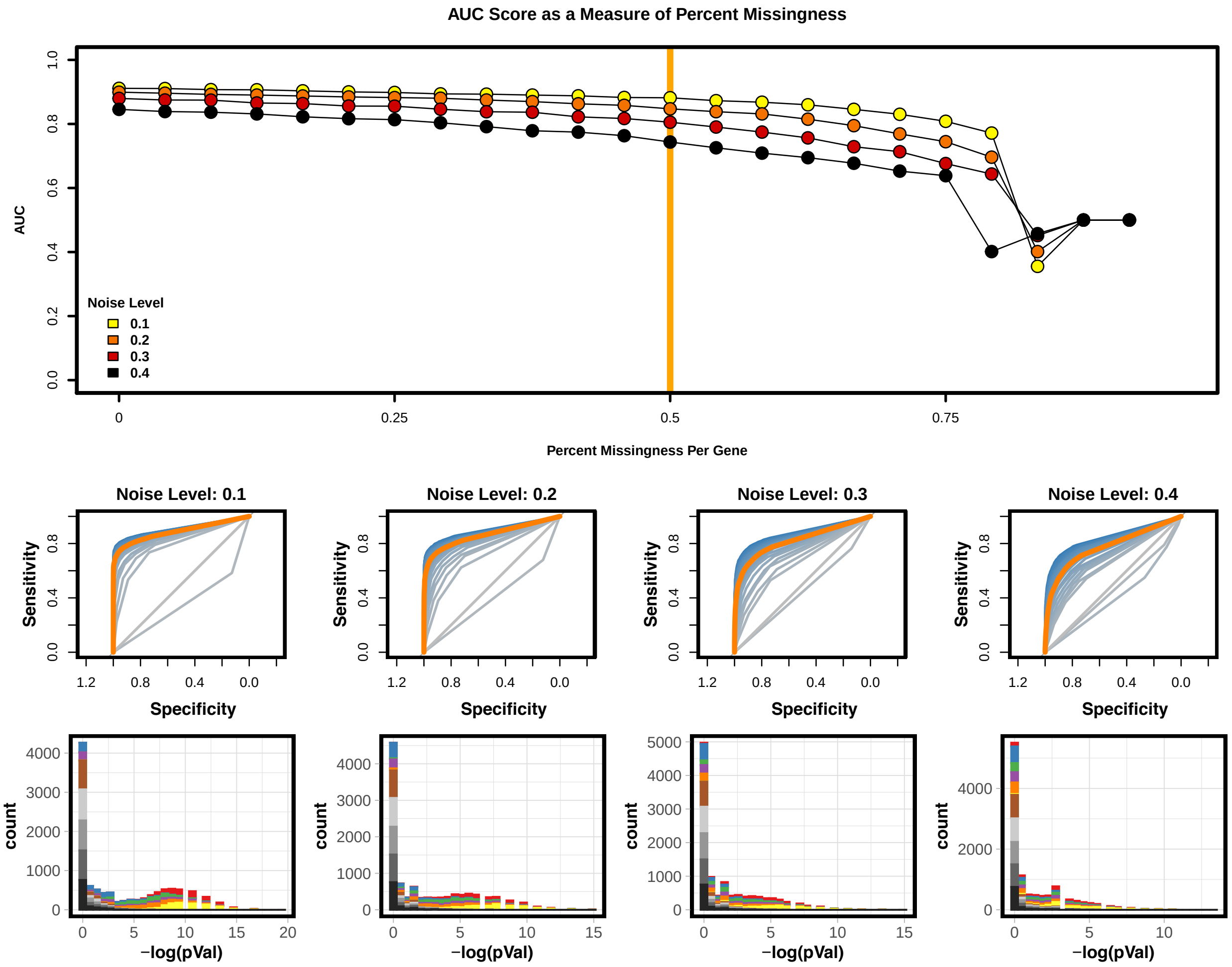


**Figure S6: Missing Data Results JTK_CYCLE | [A]** JTK_CYCLE AUC scores across different levels of percent missingness per gene for a 48-h every 2-h with 1 replicate sampling scheme with varying noise levels. 50% missingness is highlighted by the orange line. **[B] TOP:** ROC curves for each percent missingness depicted in the AUC score plot in panel A above. ROC curves scaled from Blue (0% missingness) to grey (96% missingness). 50% missingness is again highlighted by the orange curve. **BOTTOM:** Histogram of $-\log(p)$ at the different noise levels corresponding to the orange 50% missingness ROC plot.

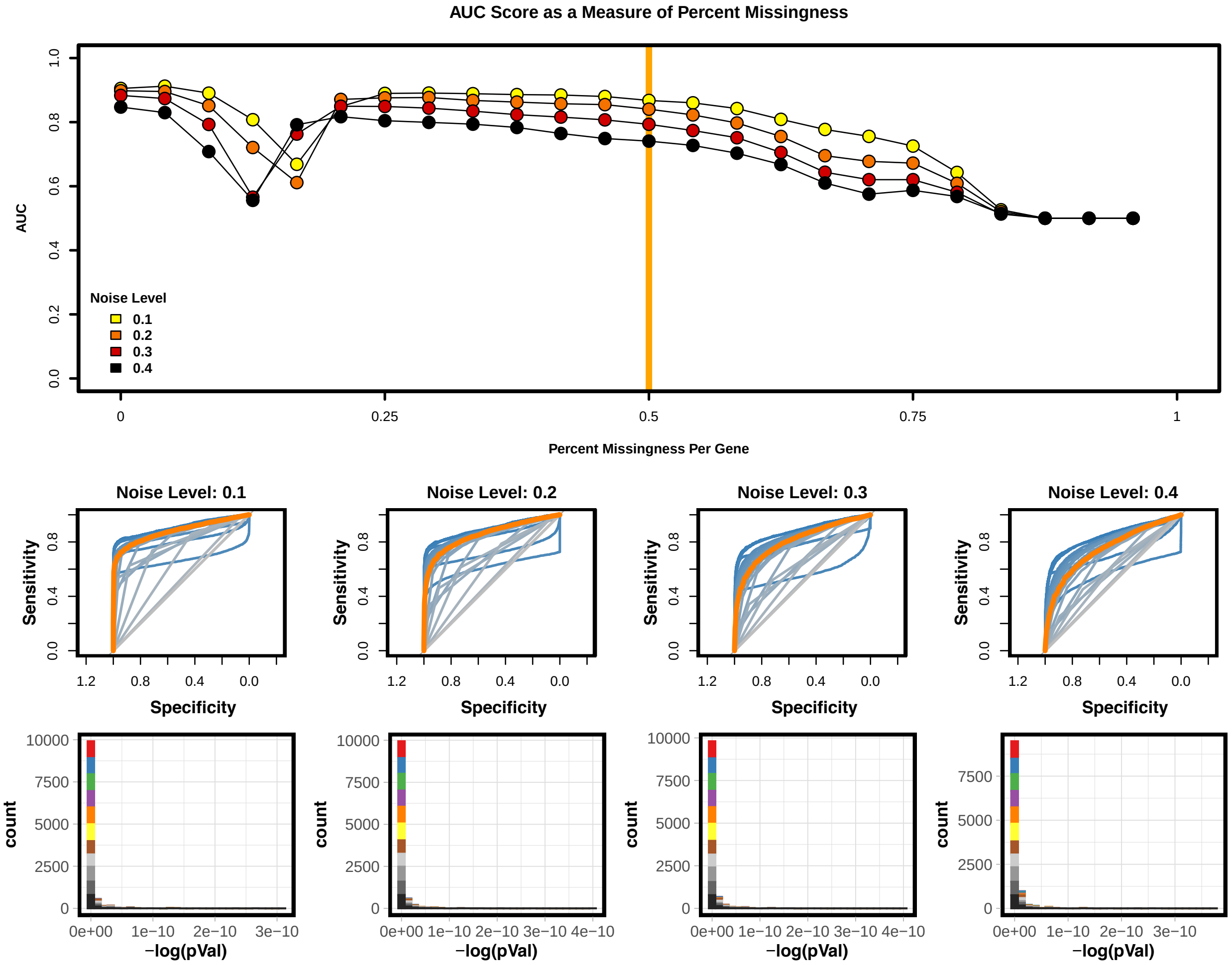


**Figure S7: Missing Data Results RAIN | [A]** RAIN AUC scores across different levels of percent missingness per gene for a 48-h every 2-h with 1 replicate sampling scheme with varying noise levels. 50% missingness is highlighted by the orange line. **[B] TOP:** ROC curves for each percent missingness depicted in the AUC score plot in panel A above. ROC curves scaled from Blue (0% missingness) to grey (96% missingness). 50% missingness is again highlighted by the orange curve. **BOTTOM:** Histogram of $-\log(p)$ at the different noise levels corresponding to the orange 50% missingness ROC plot.

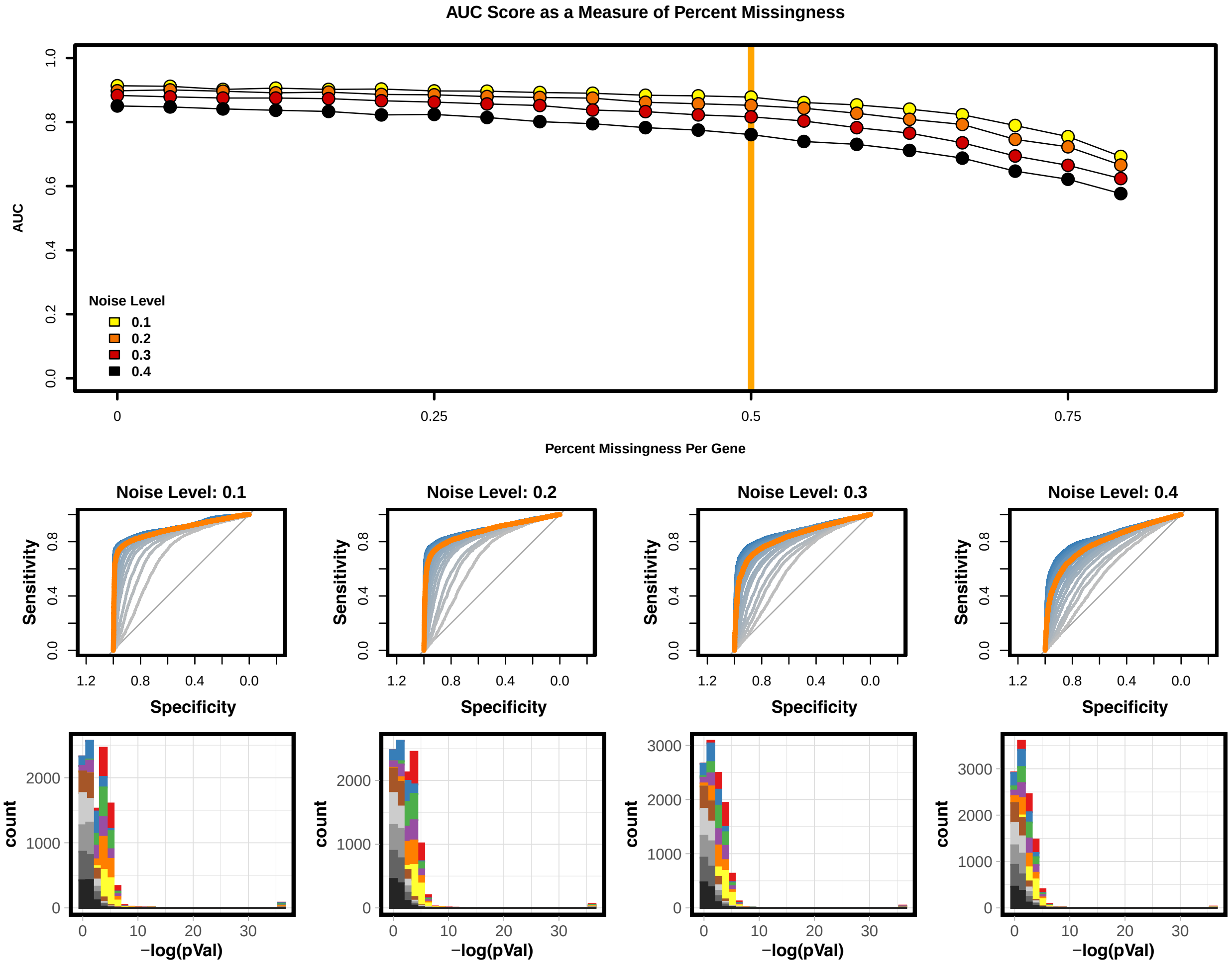


**Figure S8: Missing Data Results GeneCycle | [A]** GeneCycle AUC scores across different levels of percent missingness per gene for a 48-h every 2-h with 1 replicate sampling scheme with varying noise levels. 50% missingness is highlighted by the orange line. **[B] TOP:** ROC curves for each percent missingness depicted in the AUC score plot in panel A above. ROC curves scaled from Blue (0% missingness) to grey (96% missingness). 50% missingness is again highlighted by the orange curve. **BOTTOM:** Histogram of $-\log(p)$ at the different noise levels corresponding to the orange 50% missingness ROC plot.

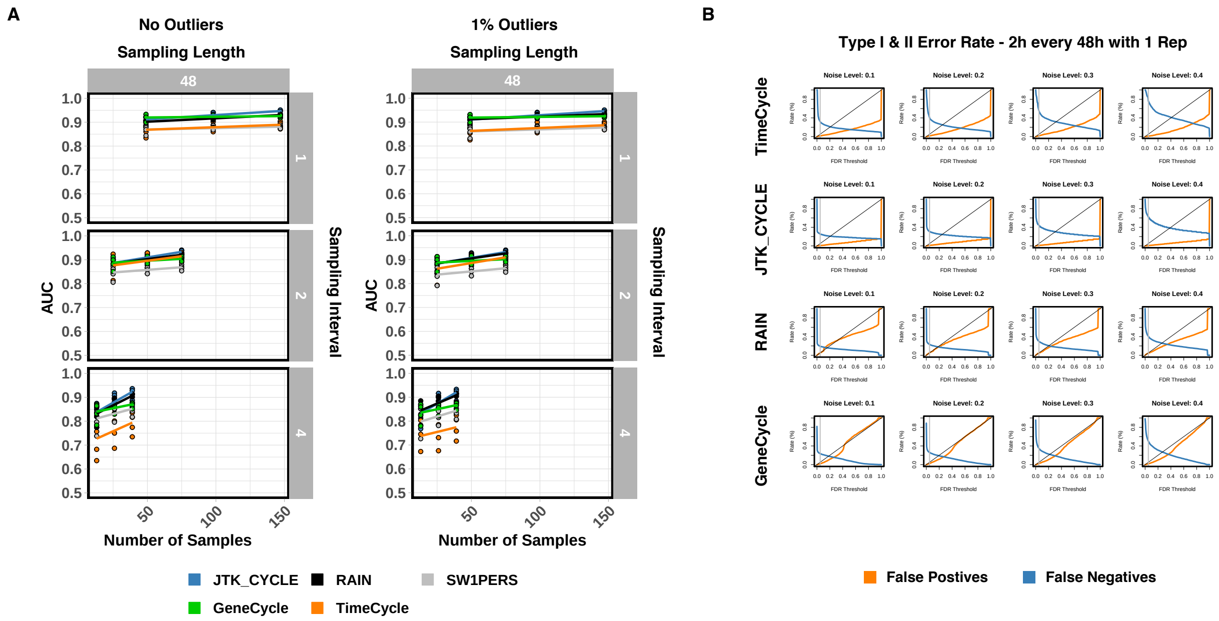


**Figure S9: Outlier Analysis** | [**A**] A total of 36 unique synthetic time-course data sets were generated in R with known ground truth, as described in [1]. Each data set consisted sampling lengths of 48-h with different number of replicates (1,2,3), sampling intervals (1-h, 2-h, 4-h), and noise levels (10%, 20%, 30%, 40%) as a percentage of the waveform amplitude. For all sampled time–points, outliers were injected into the time–series at a rate of 1%. Outliers were drawn from a uniformly distribution between $[\mu - 4\sigma, \mu - 3\sigma]$ and $[\mu + 3\sigma, \mu + 4\sigma]$ from the diurnal mean ($\mu$) for each time–series. AUC scores for each method were computed across all 36 data sets. Lines represent the best linear fit across replicates and noise levels within a specified sampling scheme. Replicate time-series were averaged together for the GeneCycle and SW1PerS algorithm, since neither algorithms has a built in method for handling replicates. Non outlier analysis plot for 48-h sampling length – identical to Figure 2A – shown for comparison. [**B**] Type I and Type II error rates at varying FDR thresholds across methods for synthetic data sampled every 2-h for 48-h with 1 replicate. FDR = 0.05 marked by grey vertical line. SW1PerS does not compute a $p$-value, but rather a periodicity score and was thus omitted from analysis.

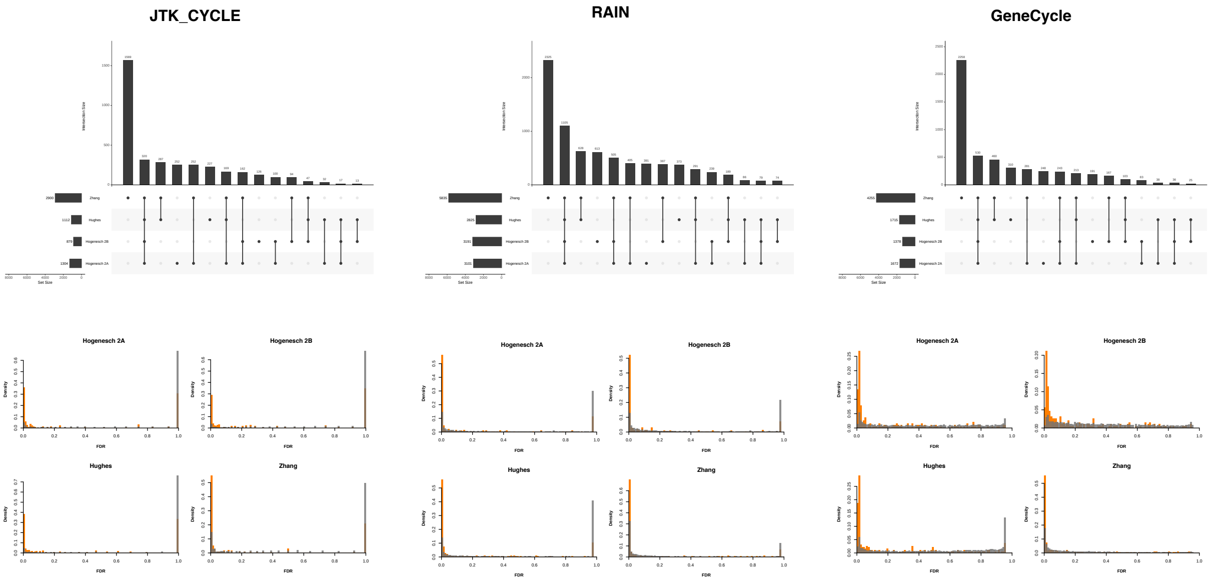


**Figure S10: Biological Data Results Methods Comparison| TOP:** Upset plot overlap between significant cycling genes sets detected by each method at an FDR $< 0.05$ across 3 distinct mouse liver time-series data sets as described in [1]. The Zhang and Hughes data sets were sampled every 2-h for 48-h. The Hogenesch study, sampled every 1-h for 48-h, was downsampled into two data sets sampled every 2-h for 48-h (Hogenesch 2A and Hogenesch 2B). **BOTTOM:** Distribution of FDR correct $p$-values of the known circadian genes (orange) versus all genes (grey) across datasets. Circadian genes were extracted from the CGDB database [2]. Only circadian genes that were experimentally validated in mouse liver tissue through low-throughput methods were included in the analysis.

### Note regarding 24-h sampling duration

It is not uncommon for researchers to elect to sample for durations of 24h for economic and practical reasons. We wish to emphasize here that this is not a suitable duration for the purposes of cycling detection. We illustrate this by applying three cycle detection methods (TimeCycle, JTK_cycle, and RAIN) to synthetic data (**Figure S11A**) and to and biological data subset to 24-h of sampling (**Figure S11B**). In **Figure S11B**, we took 24 sliding windows spanning 24-h from the 48-h Hogenesch dataset, generating 24 subsampled datasets whose only difference was the sampling start time. If a method accurately detects cycling genes, we should see a strong overlap in the majority of subsamples (i.e., genes should be called as cycling or not regardless of the sampling start time). In both the synthetic and biological data, TimeCycle classifies no gene as cycling—a logical outcome, considering that the observation of a single period is not sufficient to establish whether a signal is, in fact, periodic. Both JTK_cycle and RAIN do call some genes as cycling, and JTK_cycle appears to be accurate in the synthetic data, but the genes classified as cycling in the real data tend to only be called in one or a few of the windows, suggesting that the results are highly unreliable and depend very strongly on the sampling start time. We thus strongly caution the community against using 24-h durations for circadian research.

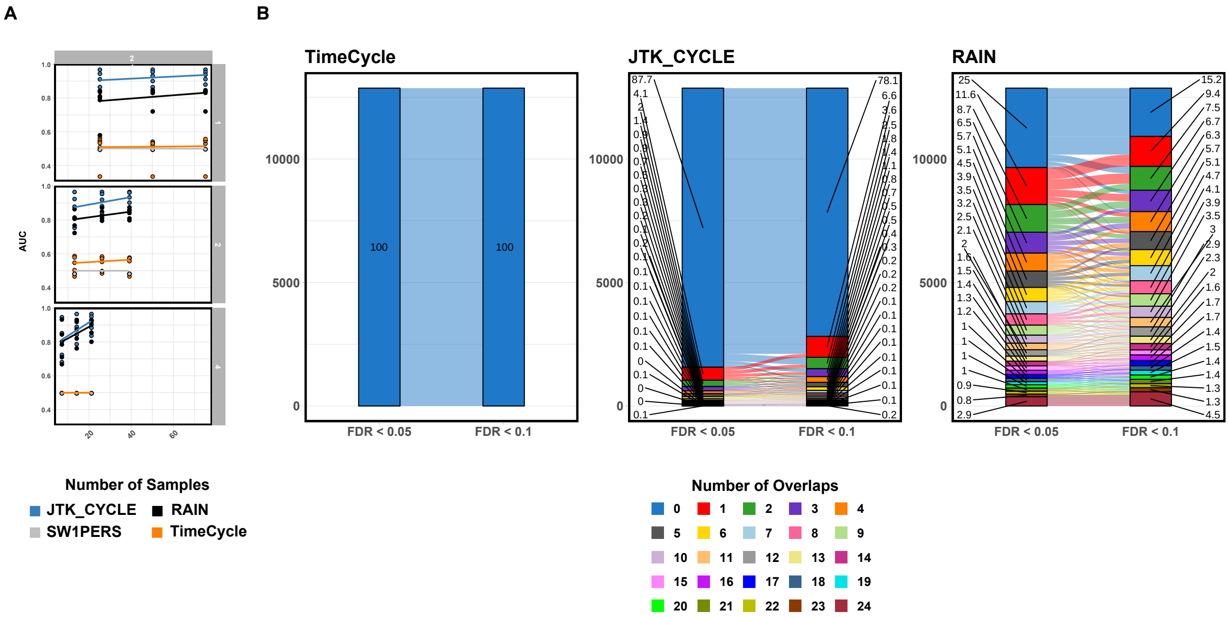


**Figure S11: 24-h Reproducibility Analysis with Synthetic and Biological Data** | **[A]** A total of 45 unique synthetic time-course data sets were generated in R with known ground truth, as described in [1]. Each data set consisted of a different number of replicates (1,2,3), sampling intervals (1-h, 2-h, 4-h), sampling lengths (24-h), and noise levels (0%, 10%, 20%, 30%, 40%) as a percentage of the waveform amplitude. AUC scores for each method were computed across all 45 data sets. Lines represent the best linear fit across replicates and noise levels within a specified sampling scheme. Replicate time-series were averaged together for the SW1PerS algorithm, since SW1PerS does not have a built in method for handling replicates. **[B]** Alluvial plot of the number of times a gene was identified as cycling in 24 sliding windows of 24-h in the Hogenesch data set. The y-axis represents raw counts with the percentages of genes in each overlap category displayed on the bar graph. Lines between bars represent the transition from the number of overlaps on a per gene basis between the sliding window frames at an FDR $< 0.05$ to an FDR $< 0.1$.

[2] Shujing Li, Ke Shui, Ying Zhang, Yongqiang Lv, Wankun Deng, Shahid Ullah, Luoying Zhang, and Yu Xue. CGDB: A database of circadian genes in eukaryotes. *Nucleic Acids Research*, 45(D1):D397–D403, jan 2017.
